# Aminoglycoside-induced premature termination codon readthrough of *COL4A5* nonsense mutations that cause Alport syndrome

**DOI:** 10.1101/2021.06.11.448099

**Authors:** Kohei Omachi, Hirofumi Kai, Michel Roberge, Jeffrey H. Miner

## Abstract

Alport syndrome (AS) is characterized by glomerular basement membrane (GBM) abnormalities leading to progressive glomerulosclerosis. Mutations in the *COL4A3, COL4A4* or *COL4A5* genes encoding type IV collagen α3α4α5 cause AS. Truncated α3, α4, and α5 chains lacking an intact COOH-terminal noncollagenous domain due to a premature termination codon (PTC) cannot assemble into heterotrimers or incorporate into the GBM. Therefore, achieving full-length protein expression is a potential therapy for AS caused by truncating nonsense mutations. Small molecule-based PTC readthrough (PTC-RT) therapy has been well studied in other genetic diseases, but whether PTC-RT is applicable to AS is unexplored. To investigate the feasibility of PTC-RT therapy in AS, we made a cDNA to express COL4A5 fused to a C-terminal NanoLuc luciferase (NLuc) to monitor full-length translation. Full-length COL4A5-NLuc produces luminescence, but mutants truncated due to a PTC do not. To screen for *COL4A5* nonsense mutants susceptible to PTC-RT, we introduced 49 individual nonsense mutations found in AS patients into the COL4A5-NLuc cDNA. Luciferase assays revealed that 11 mutations (*C29X, S36X, E130X, C1521X, R1563X, C1567X, W1594X, S1632X, R1683X, C1684X* and *K1689X*) were susceptible to PTC-RT induced by G418, which is known to have high readthrough activity. Moreover, we found that some next-generation “designer” PTC-RT drugs induced RT, and RT enhancer compounds increased the efficacy of PTC-RT in a G418-susceptible PTC mutant. These results suggest that PTC-RT therapy is a feasible approach for some patients with AS. Our luciferase-based COL4A5 translation reporter system will contribute to further development of PTC-RT therapies in a personalized medicine approach to treating AS.

## INTRODUCTION

Alport syndrome (AS) is a hereditary kidney glomerular disease with eye and inner ear defects characterized by glomerular basement membrane (GBM) abnormalities leading to progressive glomerulosclerosis and kidney failure (1). Mutations in either the *COL4A3* (2,3), *COL4A4* (4) or *COL4A5* (5) genes encoding the type IV collagen α3, α4, and α5 chains, respectively, cause AS. All three chains are necessary to form a functional type IV collagen α3α4α5 network. The type chains assemble inside cells into α3α4α5 heterotrimers (protomers), which are secreted into the extracellular space (6), where they polymerize to build a basement membrane with other components such as laminins, nidogen and heparan sulfate proteoglycans (7).

The lack or reduction of type IV collagen α3α4α5 in AS eventually leads to GBM abnormalities including thinning, thickening, and splitting. Current standard-of-care therapy uses renin-angiotensin system (RAS) inhibitors such as angiotensin-converting enzyme (ACE) inhibitors or Angiotensin II receptor blockers (ARBs). Though they delay progression to kidney failure, they do not cure AS (11-13). In contrast, development of methods to fix the pathogenic GBM abnormalities—compositional, structural, and functional—could cure AS or overcome the limitations of current treatments.

One of the potential barriers to treatment of AS using a GBM repair approach is the requirement that the abnormal GBM composed of collagen IV α1α1α2 be able to incorporate α3α4α5. Genetic rescue experiments in a *Col4a3*-null AS mouse model has shown that postnatal induction of COL4A3 production in podocytes, the glomerular cells that normally synthesize the collagen IV α3α4α5 network, enables α3α4α5 trimer synthesis, secretion, and incorporation into the Alport GBM (14). This study shows that restoration of the normal type IV collagen α3α4α5 network in the Alport GBM, by either cell-, protein-, chemical- or gene-based approach, is a feasible approach towards a cure.

In the present study, we focused on chemical-induced restoration COL4A5 expression in *COL4A5* nonsense mutation types of AS. Nonsense mutations resulting in premature termination codons (PTCs) account for about 6% of AS cases (15). Type IV collagen chains have a C-terminal NC1 domain that is essential for assembly of heterotrimers inside cells and for network formation in the GBM. Truncated α3, α4, α5 chains without an intact NC1 domain due to PTCs cannot form trimers or polymerize in the GBM (16). Therefore, achieving full-length protein expression is a potential therapy for AS due to nonsense mutations.

Small molecule-based PTC readthrough (PTC-RT) therapy has been well studied in other genetic diseases such as cystic fibrosis (17,18), Duchenne muscular dystrophy (19,20), and inherited skin disorders (21,22). G418, an aminoglycoside class antibiotic, is the most studied PTC-RT drug. Aminoglycosides bind to prokaryotic ribosomes and inhibit protein synthesis in gram-negative bacteria. In addition to their affinity for prokaryotic ribosomes, aminoglycosides are known to bind to eukaryotic ribosomes and induce PTC-RT of nonsense mutants by enabling a near-cognate aminoacyl-tRNA to recognize PTCs (23). Since the discovery of aminoglycoside-induced PTC-RT, it has been shown that PTC-RT can be induced by various aminoglycoside compounds (17,24,25). Although aminoglycoside-mediated PTC-RT has the advantages of being well-studied and highly efficient, high doses can cause nephrotoxicity and ototoxicity. To overcome these limitations, structurally designed aminoglycosides with reduced nephrotoxicity and ototoxicity that maintain readthrough activity have been developed (24,26). In addition, non-aminoglycoside PTC-RT compounds have been identified by high throughput library screening (19,27) and some have been chemically modified to improve their activity (28-30). In addition, several enhancer compounds have been found that enhance the activity of PTC-RT compounds, allowing the use of reduced doses of aminoglycosides and thus reduced toxicity (31,32).

With these technological advances, PTC-RT-based therapy has become a more realistic option. However, it is unexplored whether nonsense readthrough therapy is applicable to AS. Here, we tested the feasibility of PTC-RT therapy for nonsense mutant AS. We generated a NanoLuc-based translation reporter system to evaluate which *COL4A5* nonsense mutations are susceptible to PTC-RT therapy. 49 nonsense mutations reported in patients with AS were tested, and 11 of them were highly sensitive to aminoglycoside-mediated PTC-RT. Moreover, we found that designer aminoglycoside and non-aminoglycoside PTC-RT drugs with reduced nephrotoxicity and ototoxicity induced full-length COL4A5 protein synthesis in PTC-RT-sensitive mutants. Also, PTC-RT enhancer compounds potentiated aminoglycoside-mediated PTC-RT. These results contribute important basic knowledge for the feasibility of PTC-RT therapy in AS.

## RESULTS

### Development of a NanoLuc-based COL4A5 translation reporter system

The efficacy of aminoglycoside-induced PTC-RT varies greatly from mutation to mutation (22). At least 76 *COL4A5* nonsense mutations have been reported in patients with X-linked AS, accounting for 6% of all known X-linked AS mutations (15). Therefore, determining which nonsense mutations are susceptible to PTC-RT is a crucial first step towards clinical application. To evaluate the sensitivity of *COL4A5* nonsense mutations to PTC-RT in a high-throughput system, we generated a COL4A5-NanoLuc reporter plasmid by in-frame fusion of NanoLuc to the COOH-terminus of COL4A5 (Fig. 1A). Introduction of a PTC into the COL4A5-NanoLuc cDNA leads to synthesis of truncated protein without the COOH-terminal NanoLuc, so luminescence is not produced. In this reporter system, it is assumed that a small molecule compound such as an aminoglycoside will promote PTC-RT, leading to synthesis of a full-length protein and production of luminescence.

**Figure 1.**
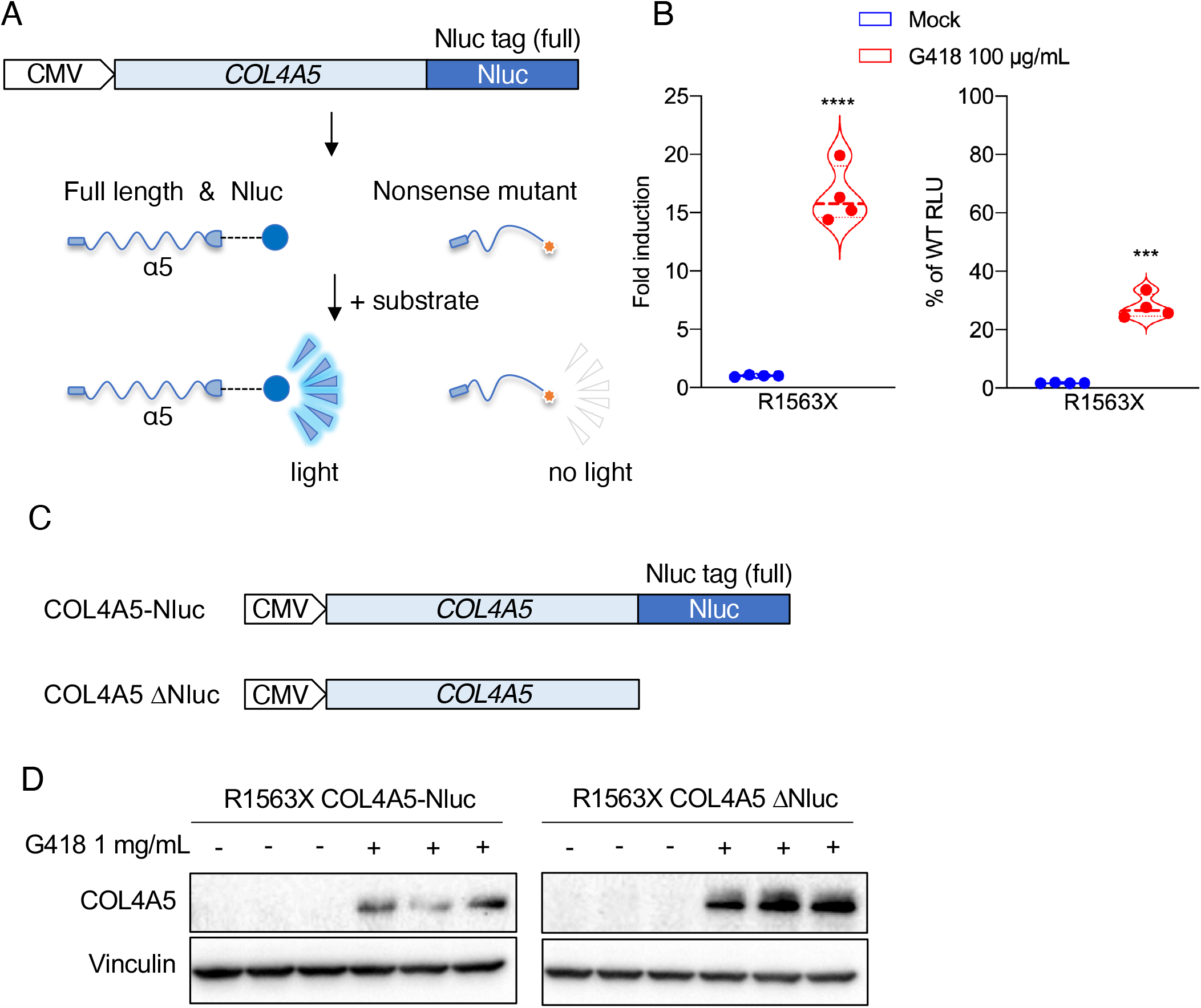
Development of a luciferase-based screening platform for testing PTC-RT of *COL4A5* nonsense mutations. *A*, Schematic representation of the NanoLuc (Nluc) luciferase-based COL4A5 PTC-RT reporter construct. Nluc was fused in-frame to the C-terminus of COL4A5. Translation of full-length COL4A5 produces a fused functional Nluc that generates luminescence, but truncation of COL4A5 translation due to a PTC results in no luminescence. *B*, Luminescence was measured in cell lysates from HEK293 cells transfected with CMV-COL4A5-WT- and -R1563X-NanoLuc plasmids and HSV-TK-Luc2 (firefly luciferase) for normalization. G418 treatment induced PTC-RT in COL4A5-R1563X-Nluc expressing cells. Statistical analysis was performed using Student’s t-test (n=4). ***, *P* <0.005; ****, *P* <0.001 versus Mock. *C*, Schematics of NanoLuc-tagged and -non-tagged COL4A5 expression constructs. *D*, Immunoblots of intracellular-Nluc-tagged or non-tagged COL4A5 products in HEK293 cells treated with G418 for 24 h. Full-length COL4A5 was detected by COL4A5 NC1 domain antibody (H52) and anti-Vinculin was used as loading control. G418 induced PTC-RT of COL4A5-R1563X in both NanoLuc-tagged and non-tagged COL4A5 expressing cells. RLU, relative light units.

First, we introduced a pathogenic nonsense mutation, R1563X, into the COL4A5-NanoLuc cDNA to see if we could detect PTC-RT in the presence of G418, which is known to have high readthrough activity and is considered the gold standard PTC-RT drug in vitro. G418 increased luminescence in HEK293 cells expressing COL4A5-R1563X-NanoLuc to a level that was 20-30% of that of WT (Fig. 1B). Moreover, to show that the PTC-RT was not artifactually related to the presence of the NanoLuc mRNA in the transcript, we investigated whether G418 could induce PTC-RT of the isolated COL4A5-R1563X mRNA, using COL4A5-R1563X-ΔNanoLuc (Fig. 1C). Immunoblot analysis showed G418 induced PTC-RT in HEK293 cells expressing either COL4A5-R1563X-NanoLuc or COL4A5-R1563X ΔNanoLuc (Fig. 1D). These results show that the COL4A5-NanoLuc reporter cDNA was sensitive and quantitative enough for monitoring translation of full-length COL4A5 protein in a multi-well plate format.

### Screening for PTC-RT-susceptible *COL4A5* nonsense mutations

To screen *COL4A5* nonsense mutations reported in patients with X-linked AS for susceptibility to PTC-RT, we introduced 49 individual mutations into the COL4A5-NanoLuc cDNA reporter by site-directed mutagenesis. Aminoglycoside-induced PTC-RT is known to be influenced by the type of PTC (UGA>UAG>UAA) (33). Therefore, we comprehensively selected *COL4A5* UGA, UAG, and UAA nonsense mutations for evaluation. Nonsense mutant COL4A5-NanoLuc plasmids were individually transfected into HEK293 cells, and the cells were treated for 24 h with different concentrations of G418. G418 induced significant PTC-RT of 40 of the 49 nonsense mutations (Fig. 2A-C). Many of them were statistically significant, but some did not have high PTC-RT rates. Of the types of mutations that responded to G418, UGA PTCs showed the highest readthrough rates, which is in line with previous studies. 11 of 49 *COL4A5* nonsense mutants (*C29X, S36X, E130X, C1521X, R1563X, C1567X, W1594X, S1632X, R1683X, C1684X, K1689X*) showed more than 5-fold induction of PTC-RT. The amount of luminescence produced from these G418-susceptible mutants was about 10-30% of the WT level (Fig. 2D).

**Figure 2.**
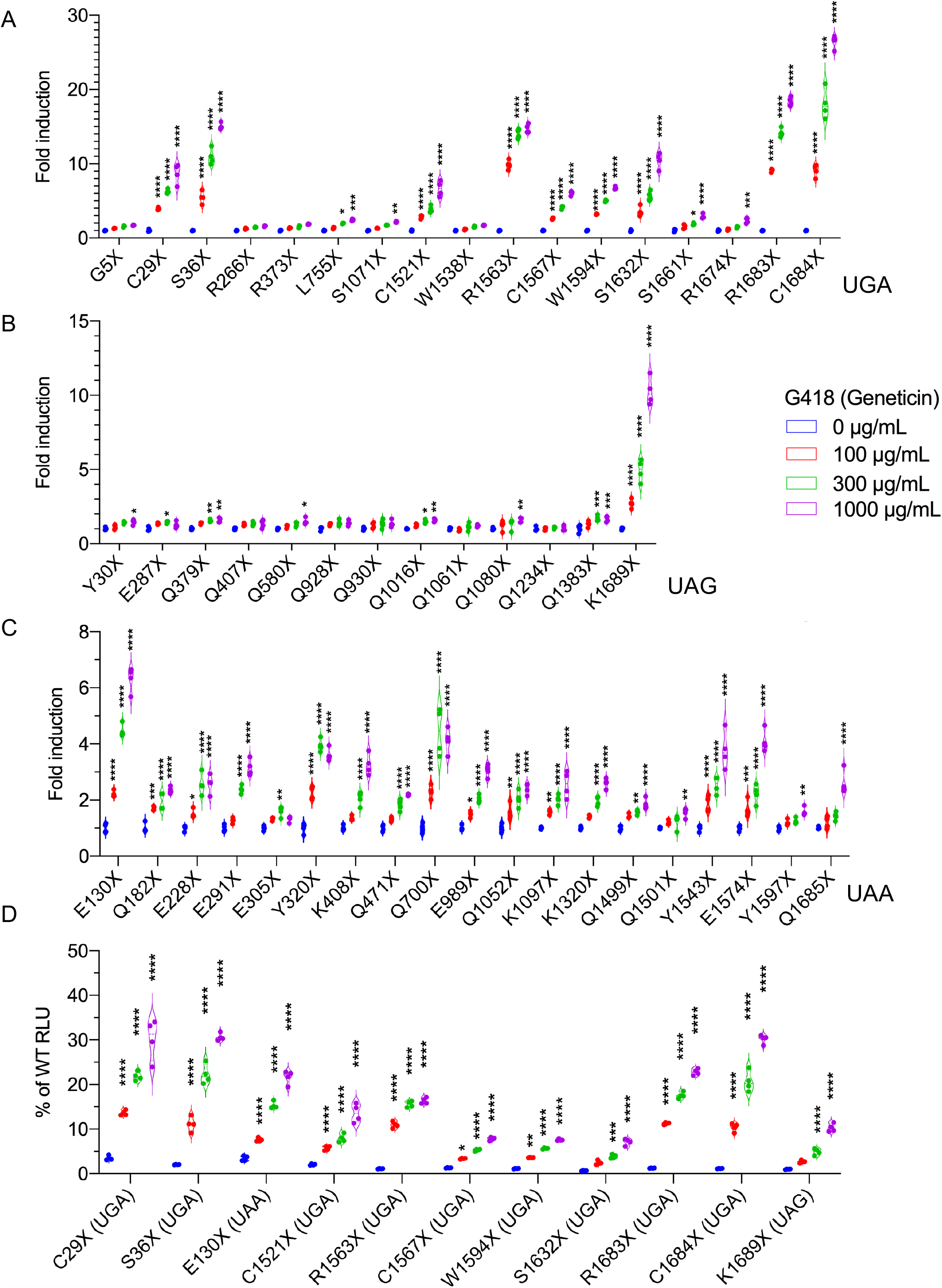
Identification of *COL4A5* mutations susceptible to G418-induced PTC-RT. *A-C*, Luminescence was measured in the cell lysates from HEK293 cells co-transfected with CMV-NanoLuc-fused COL4A5-WT or with the indicated nonsense mutants and with HSV-TK-Luc2 (firefly) for normalization. Cells expressing one UGA (*A*), UAG (*B*), or UAA (*C*) COL4A5 nonsense mutant cDNA were treated with G418 at the indicated concentrations for 24 h, and luminescence was measured. G418 induced PTC-RT of some but not all nonsense mutants. *D*, Readthrough efficiency of eleven PTC-RT-susceptible mutants was compared to WT. Statistical analysis was performed using two-way ANOVA with Tukey’s multiple comparisons test (n=4). *, *P* <0.05; **, *P* <0.01; ***, *P* <0.005; ****, *P* <0.001 vs. no treatment. RLU, relative light units.

### Gentamicin, an aminoglycoside approved for clinical use, induces PTC-RT in G418-susceptible mutants

Although G418 is one of the most potent readthrough inducers, it is highly toxic and cannot be used clinically. Gentamicin is a clinically approved aminoglycoside class antibiotic. Therefore, we investigated whether gentamicin induced PTC-RT of the G418-susceptible mutants.

COL4A5-NanoLuc reporter cDNAs with introduced nonsense mutations (*C29X, S36X, E130X, C1521X, R1563X, C1567X, W1594X, S1632X, R1683X, C1684X, K1689X*) were transfected into HEK293 cells, and the extent of PTC-RT induction by gentamicin treatment was quantified by measuring luminescence. Gentamicin significantly induced PTC-RT of G418 susceptible mutants except for *COL4A5-K1689X* (Fig. 3A). The amount of full-length protein produced with gentamicin-induced readthrough peaked at 5-10% of that of WT for most mutants (Fig. 3B).

**Figure 3.**
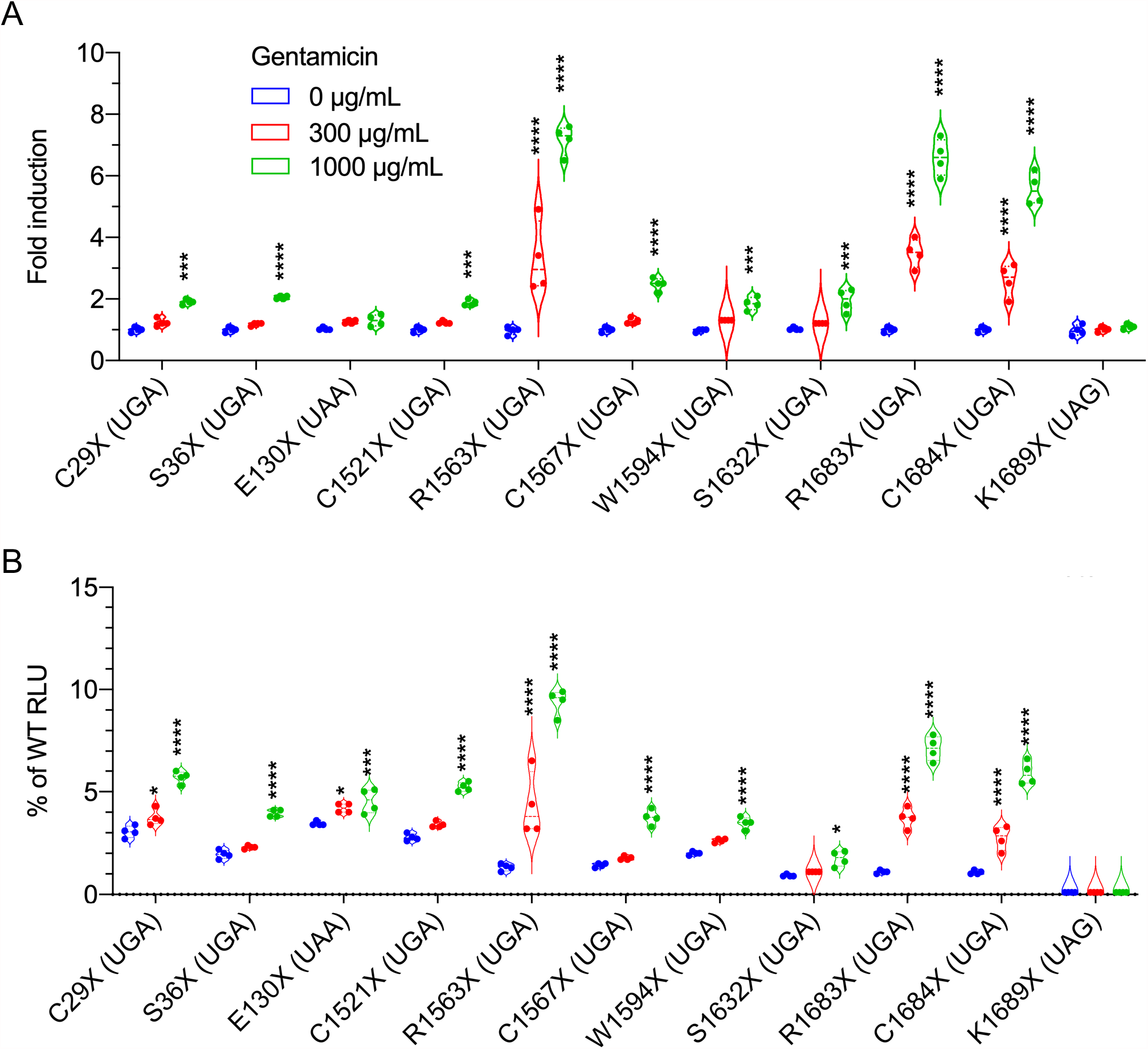
Gentamicin induced PTC-RT of G418-susceptible *COL4A5* mutations. *A*, Luminescence was measured in cell lysates from HEK293 cells co-transfected with CMV-NanoLuc-fused COL4A5-WT or with the indicated nonsense mutant plasmids and with HSV-TK-Luc2 (firefly) for normalization. COL4A5-NanoLuc expressing cells were treated with gentamicin (as indicated) for 24 h, and luminescence was measured. *B*, Readthrough efficiency of eleven PTC-RT-susceptible mutants was compared to WT. Statistical analysis was performed using two-way ANOVA with Tukey’s multiple comparisons test (n=4). *, *P* <0.05; ***, *P* <0.005; ****, *P* <0.001. vs. no treatment. RLU, relative light units

### The efficacy of aminoglycoside-mediated PTC-RT is dose- and treatment time-dependent

To investigate whether aminoglycoside-induced PTC-RT is treatment time-dependent, we performed long-term treatment experiments using low doses of aminoglycosides on cells expressing *COL4A5-R1563X* and *-R1683X* mutants that were highly responsive to G418 and gentamicin. For long-term treatment, we generated stable COL4A5-R1563X/R1683X-NanoLuc cDNA-expressing cells by lentivirus transduction. The degree of PTC-RT was increased with low-dose G418 treatment in a time-dependent manner (Fig. 4A, B). The low concentrations of G418 (10 and 30 µg/mL) slightly increased readthrough, and the moderate concentration (100 µg/mL) dramatically increased full-length protein synthesis, depending on treatment time. The longer treatment with low concentrations of gentamicin (30 and 100 µg/mL) did not increase the efficacy of PTC-RT, but readthrough was increased at the moderate (300 µg/mL) and high (1000 µg/mL) concentrations (Fig. 4C, D).

**Figure 4.**
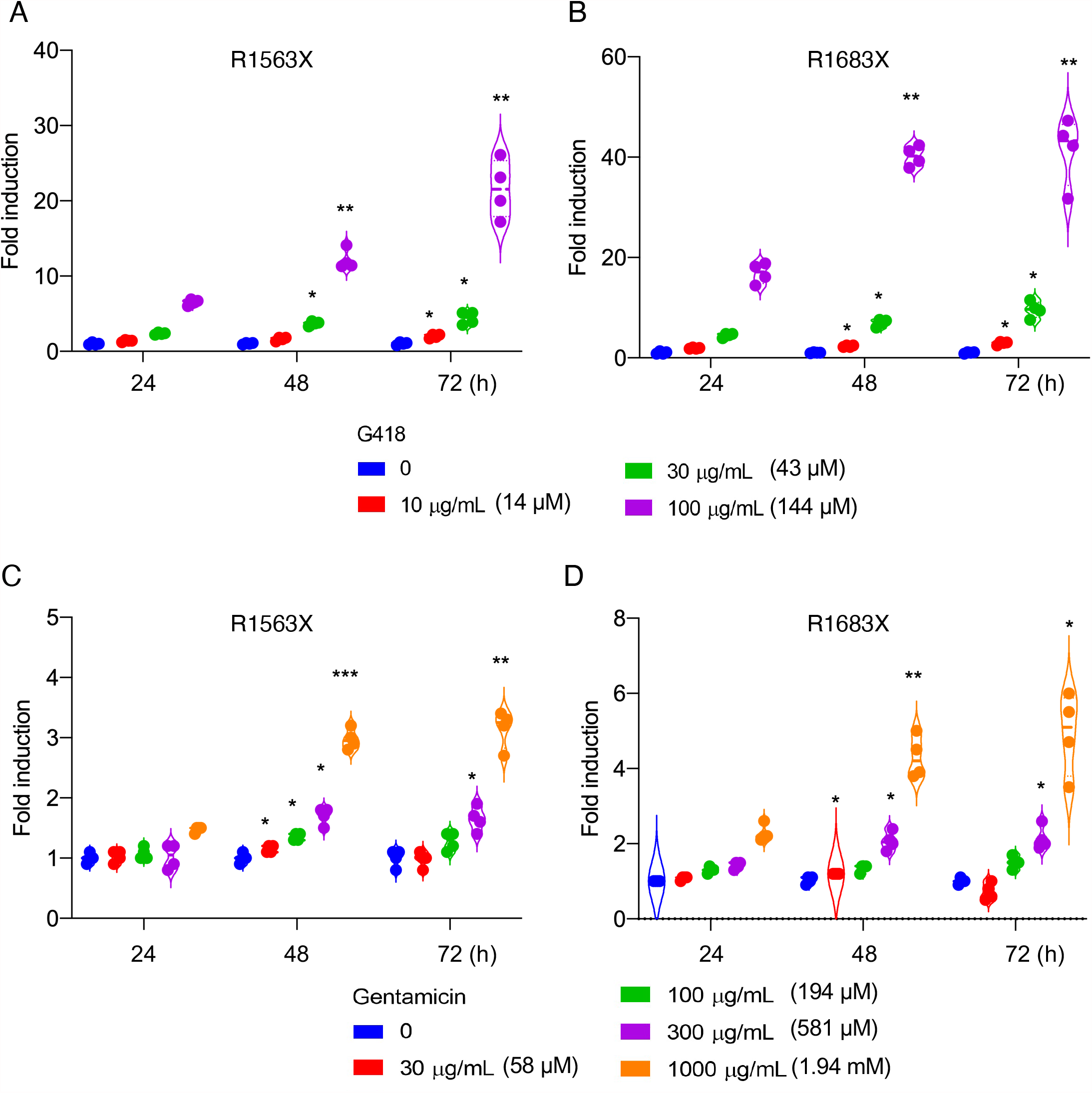
PTC-RT efficiency is dependent on both dose and treatment time. Cells stably expressing NanoLuc fused to COL4A5-R1563X or R1683X and Luc2 (for normalization) were treated with low dose G418 (*A, B*) or gentamicin (*C, D*) for the indicated times. G418 and gentamicin induced PTC-RT in the highly susceptible mutants R1563X and R1683X in a dose- and treatment time-dependent manner. Statistical analysis was performed using two-way ANOVA with Dunnett’s multiple comparisons test (n=4). *, *P* <0.05; **, *P* <0.01; ***, *P* <0.005 vs. no treatment.

### Designer aminoglycoside and non-aminoglycoside readthrough drugs induce PTC-RT in the highly susceptible mutant *COL4A5-R1563X*

The potential for successful PTC-RT therapy with aminoglycosides is dependent on their degrees of activity, nephrotoxicity, and ototoxicity (34). Aminoglycoside toxicity is attributed to a structural site different from that responsible for PTC-RT activity (24,35). Therefore, chemical modification has been used to reduce the toxicity of aminoglycosides, with the aim of reducing toxicity while maintaining readthrough activity. Several aminoglycoside derivatives have been developed and are called designer aminoglycosides (36). The use of PTC-RT compounds with non-aminoglycoside structures is also a strategy to reduce toxicity. We tested a set of next-generation PTC-RT drugs for efficacy at promoting readthrough of *COL4A5-R1563X*, a G418-susceptible mutant (Fig. 2A). HEK293 cells expressing the COL4A5-R1563X-NanoLuc cDNA were treated with several PTC-RT drugs for 24 h (Fig. 5). Whereas G418 exhibited the highest readthrough activity (Fig. 5A), ELX-02, the negamycin analogue CDX008, RTC13, and 2,6-diaminopurine (DAP) significantly induced PTC-RT dose dependently, but RTC14 and PTC124 did not (Fig. 5B-F and Fig. S1A). ELX-02 and DAP showed the highest PTC-RT activity among them (Fig. 5B, G). These results suggest that ELX-02, a designer aminoglycoside, and DAP, a purine derivative, have PTC-RT activity for G418-sensitive mutations such as R1563X, but they are expected to exhibit reduced toxicity.

**Figure 5.**
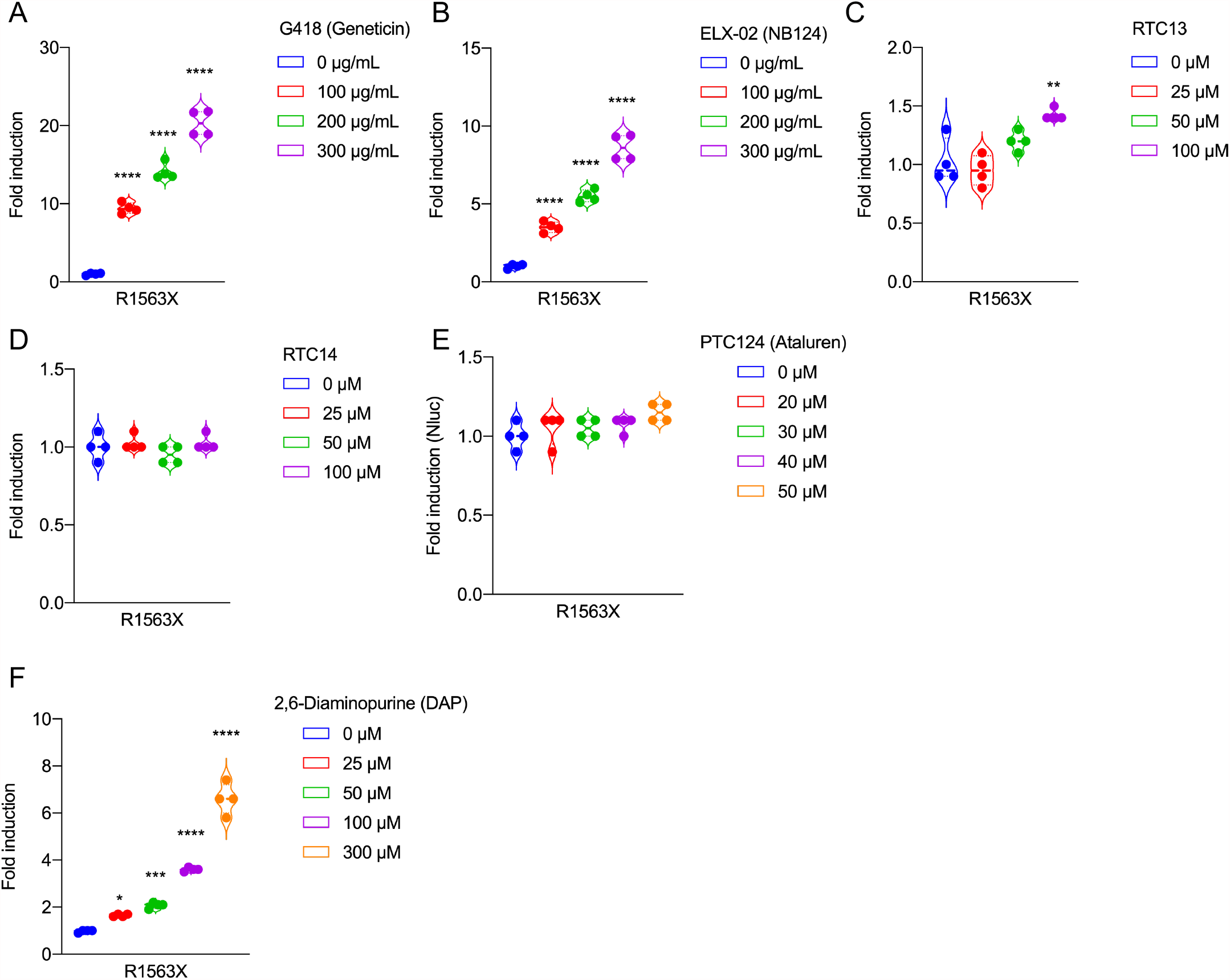
Designer aminoglycoside and non-aminoglycoside PTC-RT drugs induced PTC-RT of the highly susceptible mutant *COL4A5-R1563X*. *A-F*, Luminescence was measured in cell lysates from HEK293 cells co-transfected with CMV-NanoLuc-fused COL4A5-R1563X plasmid and HSV-TK-Luc2 (firefly) for normalization. COL4A5-R1563X-NanoLuc expressing cells were treated with serial dilutions of the indicated drugs. ELX-02, RTC13, and DAP significantly induced PTC-RT of COL4A5-R1563X. Statistical analysis was performed using one-way ANOVA with Dunnett’s multiple comparisons test (n=4). *, *P* <0.05; ***, *P* <0.005 vs. no treatment.

### Designer aminoglycoside and non-aminoglycoside PTC-RT drugs are ineffective for the non-G418-susceptible mutant *COL4A5-G5X*

In addition to the G418-susceptible *COL4A5-R1563X* mutant, we also tested whether any next generation PTC-RT drugs induced PTC-RT for the G418 non-susceptible *COL4A5-G5X* mutant (Fig. 6A). Only ELX-02 and RTC13 significantly induced PTC-RT of *COL4A5-G5X* (Fig. 6 B-F and Fig. S1B). However, the extent of induction was much less than in the case of *COL4A5-R1563X* (Fig. 5 and 6 B, D). These results indicate that PTC-RT was not strongly induced in the G418-nonsusceptible *COL4A5-G5X* mutant by either ELX-02, which has the same mechanism as G418, or DAP, which exerts its effects via a different mechanism.

**Figure 6.**
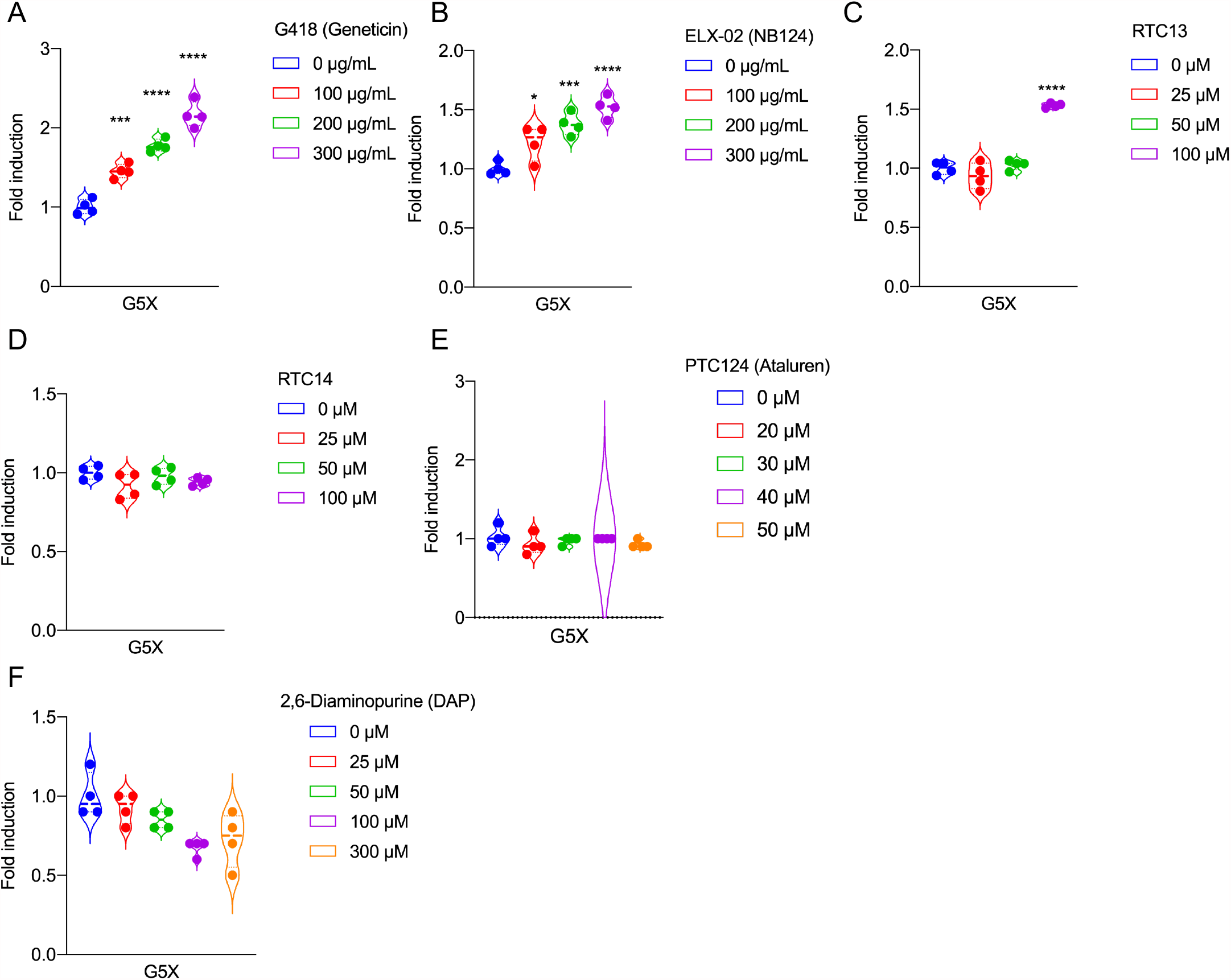
The designer PTC-RT drugs did not induce readthrough of the non-G418-susceptible mutant *COL4A5-G5X*. *A-F*, Luminescence was measured in cell lysates from HEK293 cells co-transfected with CMV-NanoLuc-fused COL4A5-G5X plasmid and HSV-TK-Luc2 (firefly) for normalization. COL4A5-G5X-NanoLuc expressing cells were treated with serial dilutions of the indicated drugs. ELX-02 and RTC13 significantly induced PTC-RT of COL4A5-G5X. However, as with the G418-mediated readthrough of COL4A5-G5X, the efficiency was lower than that of COL4A5-R1563X. Statistical analysis was performed using one-way ANOVA with Dunnett’s multiple comparisons test (n=4). *, *P* <0.05; ***, *P* <0.005 versus no treatment.

### Enhancer drugs improve the efficiency of aminoglycoside-induced PTC-RT

Several PTC-RT enhancer compounds have been developed to reduce aminoglycoside-induced toxicity by lowering the dose required for sufficient PTC-RT. The PTC-RT enhancer CDX5 was identified in a yeast cell-based assay in the presence of the aminoglycoside paromomycin. The effect of CDX5 was also significant in mammalian cells (31). A more recent study showed that the anti-malarial drug mefloquine potentiated G418-mediated PTC-RT in mammalian cells (32). Here we investigated whether readthrough enhancers potentiate aminoglycoside-mediated PTC-RT in the G418-susceptible *COL4A5-R1563X* and non-susceptible *COL4A5-G5X* mutants. Mefloquine and CDX5 derivatives potentiated both G418- and gentamicin-mediated PTC-RT of *COL4A5-R1563X* (Fig. 7A, B). On the other hand, only mefloquine potentiated both G418 and gentamicin-mediated PTC-RT of *COL4A5-G5X* (Fig. 7 C, D). Although aminoglycoside-mediated PTC-RT of *COL4A5-G5X* was enhanced by mefloquine, the induction was weaker than that for *COL4A5-R1563X* without enhancers.

**Figure 7.**
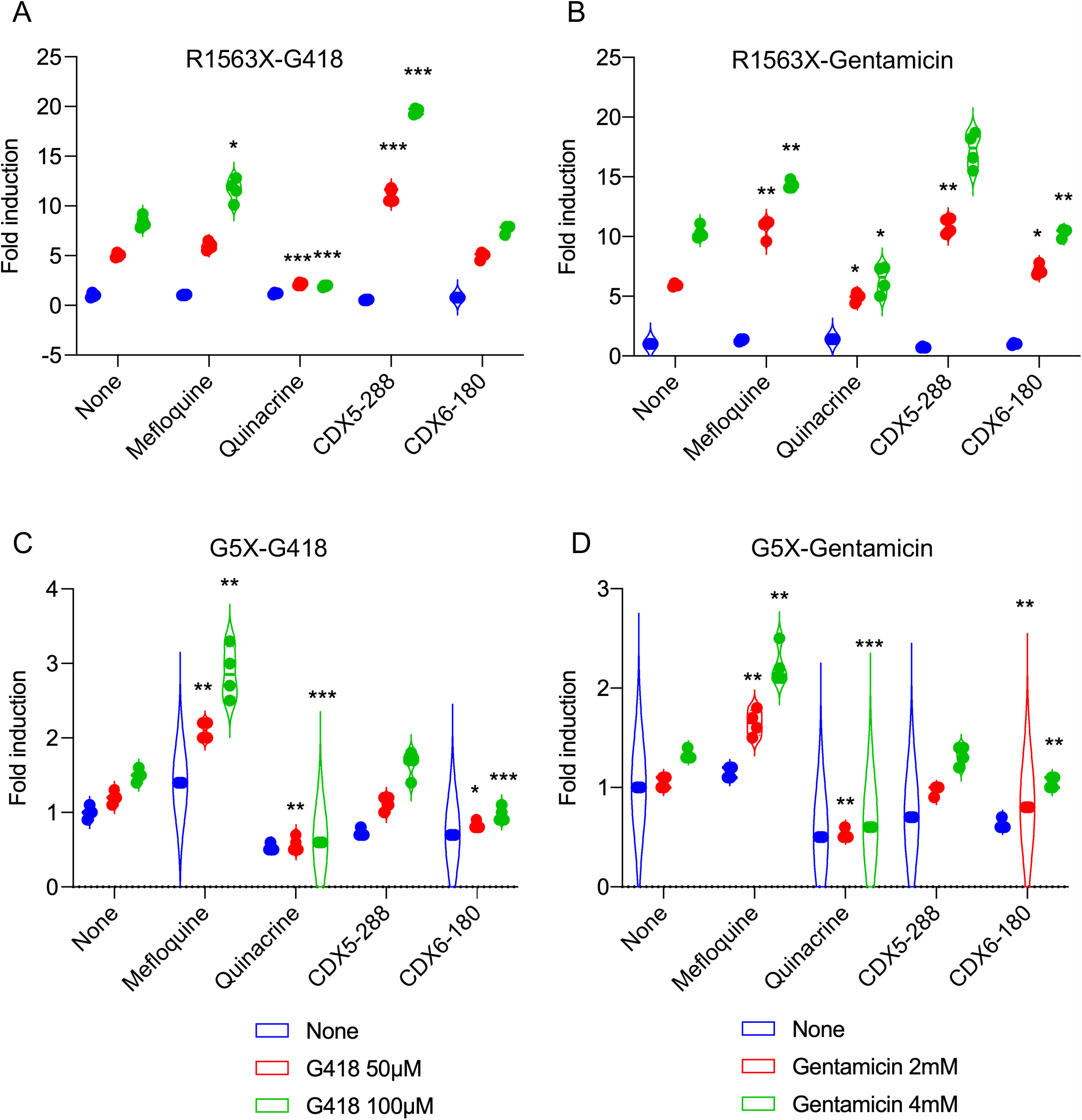
PTC-RT enhancer drugs increase the efficiency of readthrough. Luminescence was measured in the cell lysate from HEK293 cells co-transfected with CMV-NanoLuc-fused COL4A5-R1563X (*A, B*) or -G5X (*C, D*) plasmid and HSV-TK-Luc2 (firefly) for normalization. Cells were treated with the indicated doses of G418 (*A, C*) or gentamicin (*B, D*) supplemented with the indicated readthrough enhancer compounds at 20 µM. Mefloquine and CDX-288 enhanced the PTC-RT efficacy of both G418 and gentamicin in COL4A5-R1563X expressing cells. CDX6-180 slightly enhanced gentamicin-mediated PTC-RT of COL4A5-R1563X. In contrast to R1563X, only mefloquine enhanced PTC-RT of COL4A5-G5X. Statistical analysis was performed using two-way ANOVA with Dunnett’s multiple comparisons test (n=4). *, *P* <0.05; **, *P* <0.01; ***, *P* <0.005 vs. no G418 or gentamicin treatment.

### Functionality of the possible PTC-RT products of G418-susceptible mutants

So far, we have evaluated *COL4A5* nonsense mutants in terms of their susceptibility to PTC-RT, but whether the resulting protein product is functional or not is also important for therapeutic applications. PTC-RT drugs suppress PTC by facilitating the insertion of near-cognate aminoacyl-tRNAs into the ribosomal-A site during protein translation. Therefore, the readthrough product is often a full-length protein with an incorrect amino acid at the PTC. If such a substitution impairs the function of the protein, it may be difficult to rescue the mutant phenotype even if a full-length protein is produced.

To begin to investigate whether the PTC-RT products from G418-susceptible mutants are functional, we utilized a split-NanoLuc-based collagen IV α3α4α5 heterotrimer formation assay. This platform assays α3α4α5 heterotrimer formation by measuring the luminescence produced by the proximity of NanoLuc fragments fused to the ends of COL4A3 and COL4A5 that are brought together during the formation of COL4A3/4/5 heterotrimers (Fig. 8A) (40). Most pathogenic *COL4A5* missense mutations affect α3α4α5 heterotrimer formation and prevent production of functional collagen IV α3α4α5, which causes Alport syndrome. Therefore, assessing whether PTC-RT products can assemble into α3α4α5 heterotrimers is important for evaluating the feasibility of PTC-RT therapy.

**Figure 8.**
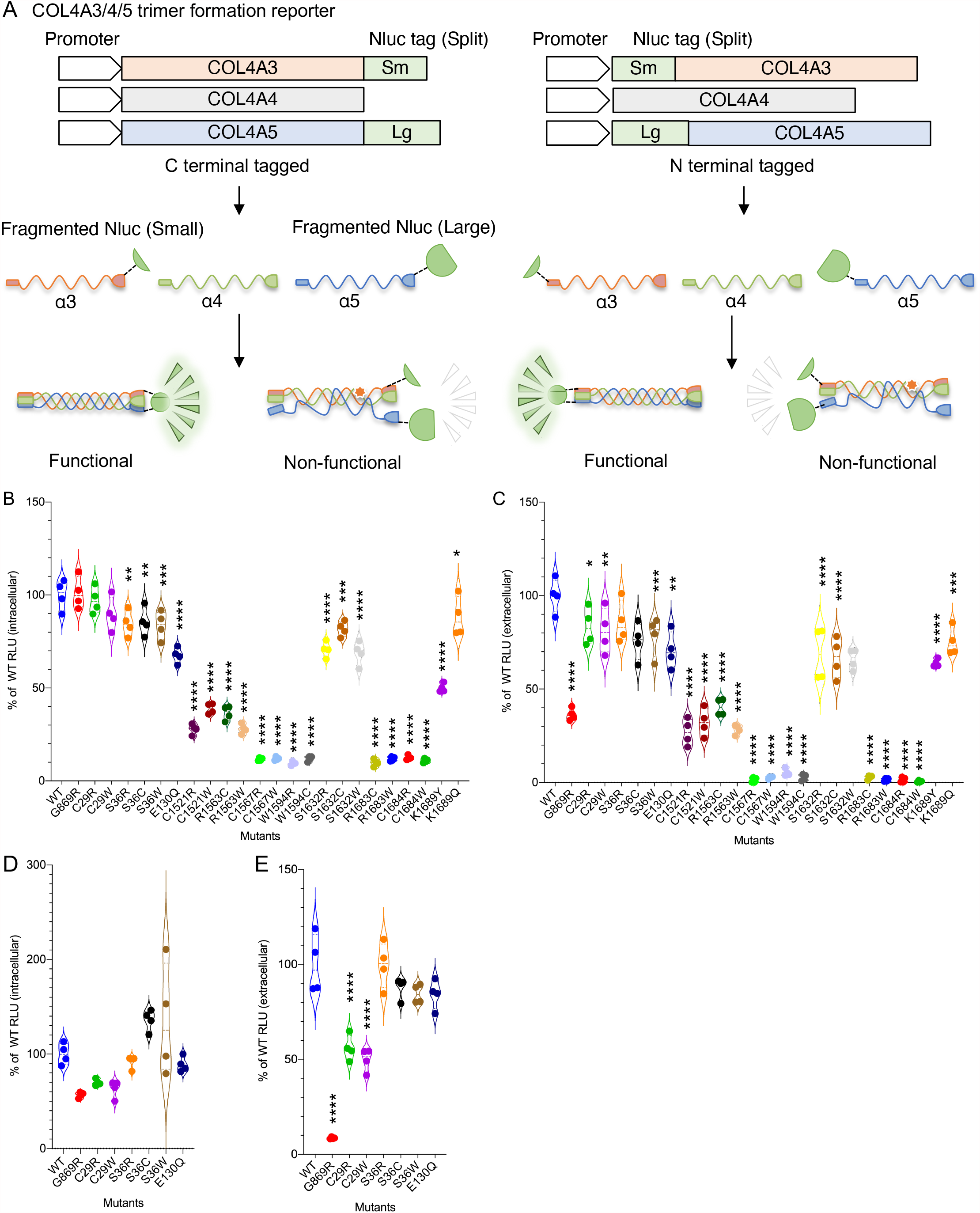
Functional analysis of the potential COL4A5 readthrough products from the PTC-RT-susceptible mutants. *A*, Schematic representation of the split NanoLuc-based COL4A3/4/5 trimer formation reporter system. Split NanoLuc fragments (large fragment: Lg, small fragment: Sm) were fused in-frame to the C- or N-termini of COL4A5 and COL4A3. When COL4A3/4/5 heterotrimers form, split NanoLuc fragments are in close proximity and acquire the ability to produce luminescence. Luminescence was measured in cell lysates (*B*) and culture media (*C*) from HEK293T cells expressing C-terminal tagged COL4A5-Lg (WT or the indicated mutants), COL4A3-Sm, and non-tagged COL4A4. The HSV-TK-Luc2 (firefly) plasmid was included for normalization. Similarly, luminescence was measured in cell lysates (*D*) and culture media (*E*) from HEK293Tcells expressing the analogous N-terminus-tagged proteins. Statistical analysis was performed using one-way ANOVA with Dunnett’s multiple comparisons test (n=4). *, *P* <0.05; **, *P* <0.01; ***, *P* <0.005; ****, *P* <0.001 vs. WT COL4A5.

It is known that during G418-induced PTC-RT, Arg, Trp and Cys are inserted for UGA, Tyr and Gln are inserted for UAA, and Gln is inserted for UAA (41). The potential readthrough products from G418-susceptible *COL4A5* mutants are shown in Table 1. In several cases, it is possible that PTC-RT will result in production of some wild-type protein. To investigate the function of the mutant readthrough products, all possible missense mutant substitutions were generated by site-directed mutagenesis and assayed using the C-terminal tagged split NanoLuc-based α3α4α5 heterotrimer assay. The luminescence reflecting heterotrimer formation was significantly decreased intracellularly and extracellularly in some readthrough products from *C1521X, R1563X, C1567X, W1594X, R1683X* and *C1684X*. On the other hand, all readthrough products from *C29X, S36X, E130X, S1632X* and *K1689X* retained the ability to form α3α4α5 heterotrimers (Fig. 8B, C). It should be noted that for Arg (R) codons mutated to UGA, more than half of the product is the wild-type R (42), so a higher percentage of functional full-length proteins are produced than for Cys (C) to UGA and Trp (W) to UGA mutants. For *C29X, S36X*, and *E130X*, the mutations are located close to the N-terminus; thus, in addition to the C-terminal tag assays (Fig. 8A, left), we also evaluated the function of appropriate PTC-RT products using the N-terminal tag system (Fig. 8A, right). The extent of heterotrimer formation for *C29X*-derived products was reduced by half, whereas *S36X-* and *E130X-*derived products retained their functions (Fig. 8D). These results indicate that inducing PTC-RT is a valid approach for a subset of COL4A5 nonsense mutations.

**Table 1.** Potential PTC readthrough products from G418-susceptible mutants.

| Nonsense mutation | PTC readthrough products | Predicted ratio (41) | Structural location |
| --- | --- | --- | --- |
| C29X (UGA) | **C29R** | 64.5 ± 11.8% | 7S domain |
|  | C29C (WT) | 17.7 ± 8.0% |  |
|  | **C29W** | 17.9 ± 6.8% |  |
| S36X (UGA) | **S36R** | 64.5 ± 11.8% |  |
|  | **S36C** | 17.7 ± 8.0% |  |
|  | **S36W** | 17.9 ± 6.8% |  |
| E130X (UAA) | **E130Y** | 47.9 ± 14.1% | Collagenous (Gly-X-Y) domain |
|  | **E130Q** | 52 ± 14.2% |  |
| C1521X | **C1521R** | 64.5 ± 11.8% | NC1 domain |
|  | C1521C (WT) | 17.7 ± 8.0% |  |
|  | **C1521W** | 17.9 ± 6.8% |  |
| R1563X | R1563R (WT) | 64.5 ± 11.8% |  |
|  | **R1563C** | 17.7 ± 8.0% |  |
|  | **R1563W** | 17.9 ± 6.8% |  |
| C1567X | **C1567R** | 64.5 ± 11.8% |  |
|  | C1567C (WT) | 17.7 ± 8.0% |  |
|  | **C1567W** | 17.9 ± 6.8% |  |
| W1594X | **W1594R** | 64.5 ± 11.8% |  |
|  | **W1594C** | 17.7 ± 8.0% |  |
|  | W1594W (WT) | 17.9 ± 6.8% |  |
| S1632X | **S1632R** | 64.5 ± 11.8% |  |
|  | **S1632C** | 17.7 ± 8.0% |  |
|  | **S1632W** | 17.9 ± 6.8% |  |
| R1683X | R1683R (WT) | 64.5 ± 11.8% |  |
|  | **R1683C** | 17.7 ± 8.0% |  |
|  | **R1683W** | 17.9 ± 6.8% |  |
| C1684X | **C1684R** | 64.5 ± 11.8% |  |
|  | C1684C (WT) | 17.7 ± 8.0% |  |
|  | **C1684W** | 17.9 ± 6.8% |  |
| K1689X | **K1689Y** | 0.8 ± 7.0% |  |
|  | **K1689Q** | 86.5 ± 8.3% |  |

## DISCUSSION

The goal of this study was to determine the applicability of PTC-RT therapy in Alport syndrome caused by nonsense mutation. Since the susceptibility to PTC-RT varies greatly among mutations (43,44), the first step would be to determine which mutations are susceptible. Because the type IV collagen genes are relatively large with many reported nonsense mutations (15,45), we thought it would be essential to evaluate a simple reporter system with high throughput to cover most of them. In addition, previous studies have shown that PTC-RT activity is affected by the sequences surrounding the nonsense mutations (33,46). Therefore, we constructed a reporter using the full-length *COL4A5* cDNA instead of a short cDNA containing the PTC. Since the full-length COL4A5 cDNA itself is about 5 kb, we used NanoLuc as a reporter to construct the fusion gene because of its small size (513 bp) and high sensitivity (47,48). The COL4A5-NanoLuc reporter cDNA developed in this study allowed us to identify which of 49 tested *COL4A5* nonsense mutations susceptible to PTC-RT. Also, this study showed the efficacy of next generation PTC-RT drugs and potentiator compounds that enhance PTC-RT activity.

Through comprehensive mutation screening, we found that UGA-type *COL4A5* nonsense mutations were more susceptible to aminoglycoside-mediated PTC-RT than other stop codons. This result is consistent with previous reports (43). However, as previously reported, not all UGA PTCs showed high susceptibility (49), and it was reconfirmed that susceptibility varied depending on the mutation. Susceptibility is known to be affected by the surrounding nucleotide sequence. To attempt to determine whether susceptibility is based on the flanking nucleotide sequence, we aligned the cDNA sequences around PTCs that were G418-susceptible and compared them to those flanking the non-susceptible PTCs (Fig. S2). However, no overt differences between susceptible and non-susceptible sequences were observed. This suggests that susceptibility is defined by factors other than the peripheral sequence, such as the location of the mutation in the gene. Although more detailed comparative studies are needed, these results highlight the importance of screening for PTC-RT using full-length cDNAs rather than just short cDNA reporters carrying sequence peripheral to the PTC.

The limitation of this reporter system is that the presence of nonsense-mediated mRNA decay (NMD) cannot be taken into account because the cDNA is already spliced and does not have the exon-exon junctions that are required for mRNA surveillance (50). The mRNAs produced from genes with nonsense mutations are partially degraded by NMD (51). Therefore, promotion of basal-readthrough by suppression of NMD is one of the therapeutic strategies for nonsense mutations, but we have not been able to investigate this. However, since the induction of PTC-RT by NMD inhibition alone is not expected to be very high (52), and many aminoglycosides have an activity that inhibits NMD itself, this limitation is not likely a serious problem, but it should be taken into account to accurately determine PTC-RT activity in vivo. To overcome this limitation, CRISPR/Cas9-mediated genome editing could be used to create cells with point mutations in endogenous genes (53), which would allow the evaluation of PTC-RT activity under the same conditions as *in vivo*. However, the throughput of this method would be very low, and it is not suitable for evaluating a large number of mutants as in the present study.

In addition to defining the readthrough susceptibility of each mutation, it is also important to determine whether the possible PTC-RT products are functional (54). To investigate whether readthrough products from G418-susceptible mutants are functional, we used split-NanoLuc-based type IV collagen α3α4α5 heterotrimer formation assays. This assay allowed us to identify which full-length but missense mutant proteins are functional. Some mutations had high readthrough activity but may lose function when a different amino acid from the original is inserted. Many of the mutations are located in the COL4A5 C-terminal NC1 domain, which is essential for α5 to form a functional triple-helical structure with other α-chains (16). Therefore, structural changes in the protein due to missense mutations are likely to interfere with the formation of the correct NC1 complex. However, the results showed that the wild-type S1632 and K1689 residues are not required to form the NC1 complex (Fig. 8 B, C). Also, the C29, S36 and E130 residues in the N-terminal 7S domain could be replaced without a total inhibition of heterotrimer assembly (Fig. 8 B-E). The results suggested that C29 substitution did not impact NC1 complex formation, but partially affected assembly at the N-terminus. This is consistent with the importance of the Cys residue in the 7S domain for disulfide bond formation to other α-chains (55). Using two different assay systems, the PTC-RT reporter assay and the α3α4α5 heterotrimer assay, we identified several mutations that seem truly susceptible to PTC-RT therapy.

The biggest challenge in PTC-RT therapy is the toxicity of drugs used at high concentrations for long periods of time to induce synthesis of enough full-length proteins to impact phenotypes. Treatment with high concentrations of aminoglycosides involves the risk of nephrotoxicity and ototoxicity. Fortunately, these issues are being addressed by the development of new PTC-RT drugs, including new aminoglycoside derivatives (24,25) and non-aminoglycoside compounds (27,30,38). Regarding this point, we showed an aminoglycoside derivative and non-aminoglycoside PTC-RT drugs induced PTC-RT in the G418-susceptible *COL4A5-R1563X* mutant (Fig. 5). In addition, from the viewpoint specific to Alport syndrome, type IV collagen α3α4α5 incorporated into the GBM should be stable for a long period of time (56). This means that once enough type IV collagen α3α4α5 is induced by PTC-RT and incorporated into the GBM, continuous treatment should not be necessary, though intermittent treatments would likely be required. This suggests that Alport syndrome is especially suitable for PTC-RT therapy.

In summary, the present study proposes PTC-RT as a novel personalized therapeutic approach for Alport syndrome based on the susceptibility of specific pathogenic nonsense mutations to readthrough. Forms of AS caused by nonsense mutations, which are classified as truncating mutations, are typically more severe than the non-truncating forms, which are usually caused by missense mutations. Therefore, the successful development of PTC-RT therapy would have significant benefits for patients with the most severe forms of AS. With various innovations such as the development of designer aminoglycosides and non-aminoglycoside PTC-RT compounds, PTC-RT therapy has become an increasingly realistic approach. In fact, some are undergoing clinical trials for nephropathic cystinosis (ClinicalTrials.gov Identifier: NCT04069260) and cystic fibrosis (ClinicalTrials.gov Identifier: NCT04126473, NCT02139306). The present study provides important information on PTC-RT-susceptible *COL4A5* mutants, and it is hoped that new gene-edited mouse models of Alport syndrome carrying the analogous mutations in *Col4a5* will facilitate proof-of-concept PTC-RT studies *in vivo* in the near future.

### Experimental procedures

#### Chemical compounds

G418 disulfate solution (50 mg/mL), RTC13, and DAP were purchased from Sigma-Aldrich (catalog no. G8168, SML1725, and 247847). Gentamicin (50 mg / mL) was purchased from Gibco, Life Technologies Corporation (catalog no. 15750-060). ELX-02 was synthesized by Sussex Research. The negamycin analog CDX008 (Fig. S1) was synthesized by WuXi AppTec. RTC14 was from ChemBridge Corporation (catalog no. 5311257). PTC124 was purchased from Cayman Chemical (catalog no. 16758).

#### Plasmids

To generate the COL4A5 with C-terminal NanoLuc fusion expression vector, full-length human COL4A5 cDNA was amplified from pEF6-COL4A5-Myc (40), cloned into pNLF1-C [CMV/Hygro] vector (Promega). pLV-BSD COL4A5-LgBiT (C terminal tag), pLV-Hygro COL4A3-SmBiT (C terminal tag), pLV-Puro COL4A4, pFN33K-COL4A5-LgBiT (N terminal tag) and pFN35K-COL4A3-SmBiT (N terminal tag) were used for split NanoLuc luciferase-based COL4A3/4/5 trimer formation assay (40). For all luciferase assays, pGL4.54 [luc2/TK] (Promega) was used as a co-transfected control vector. The mutant COL4A5 expression vectors used in this study were generated by site-directed mutagenesis as previously described. Primer sequences are shown in Table S1. The introduced mutations were verified by Sanger sequencing.

#### Cell culture and Cell lines

Human embryonic kidney (HEK) 293 cells (ATCC CRL-1573) and 293T cells (ATCC CRL-3216) cells were maintained at 37 °C, 5% CO_2_ in Dulbecco’s Modified Eagle’s Medium (DMEM) supplemented with 10% heat inactivated fetal bovine serum (FBS) and penicillin-streptomycin. For PTC-RT experiments, HEK293 cells were used because 293T cells are G418-resistant. For function tests of the potential PTC-RT products by split NanoLuc assay, 293T cells were used. Cells stably expressing cDNAs were generated by lentivirus infection. Transduced cells were selected by culturing in DMEM with appropriate antibiotics for 2 weeks.

#### Transfection, Lentivirus production, Infection and Treatment

HEK293 cells were transfected with pNLF1-C-COL4A5-Nluc (WT and mutants) and pGL4.54 [luc2/TK] plasmids by FuGENE 6 transfection reagent (Promega). Formation of plasmids/FuGENE 6 complexes was performed according to the manufacturer’s instructions. At 48 h after transfection, cells were treated with DMEM containing G418 or other PTC-RT drugs.

To produce lentivirus, 293T packaging cells were seeded at 5.5-6.0×10^5^ cells per wells in DMEM in 6-well tissue culture plates. Seeded cells were incubated at 37 °C, 5% CO_2_ for ∼20 h. Culture media were changed to fresh DMEM with 10% FBS and transfected with 1 µg of psPAX2 (Addgene: #12260), 100 ng of pMD2.G (Addgene: #12259), and 1 µg of lentivirus transfer vector per well by Lipofectamine 3000 transfection reagent (Invitrogen) according to the manufacturer’s instructions. 24 h after transfection, culture media were changed to DMEM supplemented with 30% FBS. Lentivirus-containing supernatants were collected after 24 h and filtered with 0.45 µm PVDF or PES membrane syringe filter unit. HEK293 cells or 293T cells were seeded in filtered lentivirus containing media supplemented with 0.8 µg/mL polybrene (Sigma) and cultured for 24 h, then cells were cultured in DMEM with 10% FBS and the appropriate antibiotics for 2 weeks (Hygromycin; 200-400 µg/mL, Blasticidin: 10 µg/mL, Puromycin 10 µg/mL).

#### Cell Lysis, Gel Electrophoresis, and Immunoblotting

Transfected HEK293 cells were washed twice with ice-cold PBS and lysed in RIPA buffer (0.05 M Tris-HCl [pH 7.5], 0.15 M NaCl, 1% v/v Nonidet P-40, 1% w/v Na deoxycholate, and 1% protease inhibitor cocktail). The cell lysates were centrifuged at 14,000 g for 15 min at 4 °C, and clear supernatants were collected. The protein concentration was determined using a bicinchoninic acid kit (Thermo), and equal amounts of protein lysates were loaded and separated by SDS PAGE, immunoblotted with anti-COL4A5 NC1 antibody (H52, Chondrex) and anti-vinculin antibody (7F9, Santa Cruz), and visualized using SuperSignal West Pico Chemiluminescent Substrate (Thermo).

#### Luciferase Assay

pNLF1-C-COL4A5-NanoLuc and HSV-TK-Luc2 plasmids were transfected into HEK293 cells. After 48 h, culture media were changed to culture media containing the test compounds. At 72 h after transfection, Nano-Glo Dual Luciferase Reporter Assay reagent (Promega) was added, and the luciferase activity in the cell lysates was measured using a GloMax Navigator system (Promega). All luciferase assays were conducted in LumiNunc 96-well white plates (Invitrogen). NanoLuc luciferase was normalized by constitutively expressed firefly luciferase.

#### Statistical analysis

Statistical parameters are reported in the Fig. Legends. Immunoblot experiments were performed in triplicate using 3 independent transfections. Luciferase assays were performed in quadruplicate using 4 independent cell cultures. The significance of differences between two groups was assessed using Student’s unpaired two-tailed t-tests. For three-group comparisons, we used analysis of variance (ANOVA) with Tukey-Kramer post-hoc or Dunnett’s tests. Differences with *P* values of less than 0.05 were considered statistically significant.

## Data availability

All data are presented in the main manuscript and supporting information.

## Acknowledgements

We thank David Powell of Chinook Therapeutics for synthesis and validation of ELX-02 and CDX-008.

## Author Contribution

K.O. designed the research, conducted experiments, and wrote the manuscript. J.H.M. H.K. and M.R. designed the research and edited the manuscript. All authors discussed the results and provided input on the manuscript.

## Funding and Additional Information

This work was supported by a grant from the National Institutes of Health (R01DK078314 to J.H.M.), the Children’s Discovery Institute of Washington University and St. Louis Children’s Hospital (MI-II-2019-796 to J.H.M.), the Japan Society for the Promotion of Science (JSPS) (JP19H03379 to H.K.), the JSPS Program for Advancing Strategic International Networks to Accelerate the Circulation of Talented Researchers (S2803 to H.K.), and the JSPS Program for Postdoctoral Fellowships for Research Abroad (to K.O.).

## Conflict of interest

H.K. holds a patent related to this work, Japanese Patent Application No. 2017-99497. J.H.M. receives research support from Chinook Therapeutics and the Alport Syndrome Foundation. The other authors report no conflicts of interest.

## Supplementary Material

This article contains supporting information.

## Abbreviations

The abbreviations used are

AS: Alport syndrome
PTC: premature termination codon
PTC-RT: PTC readthrough
COL4A3/A4/A5: collagen IV α3/4/5
GBM: glomerular basement membrane
luc: luciferase
NMD: nonsense-mediated mRNA decay
ANOVA: analysis of variance

## Notes

### Competing Interest Statement

H. Kai holds a patent related to this work, Japanese Patent Application No. 2017-99497. J.H. Miner receives research support unrelated to this work from Chinook Therapeutics and the Alport Syndrome Foundation. The other authors report no competing interests.

