## supporting information for "Aminoglycoside-induced premature termination codon readthrough of *COL4A5* nonsense mutations that cause Alport syndrome"

**Contents:**

Table S1. Primers used for site-directed mutagenesis of COL4A5

Figure S1. Negamycin analogue CDX008 induces PTC-RT in the G418 susceptible COL4A5-R1563X mutant

Figure S2. cDNA sequence surrounding PTCs in G418-susceptible and non-susceptible mutants

Table S1. Primers used for site-directed mutagenesis of COL4A5

| Mutation | Sense (5' to 3') | Antisense (5' to 3') |
|---|---|---|
| G5X | atgaaactgcgtTgagtcagcctggctgc | gcagccaggctgactcAacgcagtttcat |
| C29X | gcagaggctgcggcttgAtatgggtgttctccagg | cctggagaacacccataTcaagccgcagcctctgc |
| Y30X | gaggctgcggcttgctaGgggtgttctccaggatc | gatcctggagaacacccCtagcaagccgcagcctc |
| S36X | ctatgggtgttctccaggatGaaagtgtgactgcagtggc | gccactgcagtcacactttCatcctggagaacacccatag |
| E130X | gatgcaatggaaccaagggaTaacgtggatttccaggcag | ctgcctggaaatccacgttAtcccttggttccattgcatc |
| Q182X | tatccaggtcctcctggaataTaaggcctacctggtcccac | gtgggaccaggtaggccttAtattccaggaggacctggata |
| E228X | aaatttccagggacccaaaggtTaaaaaggtgagcaaggtcttcag | ctgaagaccttgctcacctttttAacctttgggtccctggaaattt |
| R226X | gacttcctggtgacTgagggcctcctgga | tccaggaggccctcAgtcaccaggaagtc |
| E287X | ccaggtggtgagaaaggtTagaagggtgagcaaggagag | ctctccttgctcacccttctAacctttctcaccacctgg |
| E291X | gaaaggtgagaagggtgagTaaggagagccaggcaaaagagg | cctcttttgcctggctctccttActcacccttctcacctttc |
| E305X | ggtaaaccaggcaaagatggaTaaaatggccaaccaggaattc | gaattcctggttggccattttAtccatctttgcctggtttacc |
| Y320X | tgcctggtgatcctggttaAcctggtgaacccggaagggatg | catcccttccgggttcaccaggTtaaccaggatcaccaggca |
| R373X | ggagaaaaaggagagTgaggatttcctggaataca | tgtattccaggaaatcctcActctcctttttctcc |
| Q379X | gcgaggatttcctggaataTagggtccacctggcctt | aaggccaggtggaccctAtattccaggaaatcctcgc |
| Q407X | cctggagaaaggggtTagaaaggtgatgaaggaccac | gtggtccttcatcacctttctAaccccttctctccagg |
| K408X | cctggagaaaggggtcagTaaggtgatgaaggaccacct | aggtggtccttcatcaccttActgacccctttctccagg |
| Q471X | gataaaggactccaaggagaaTaaggagtgaaaggtgaca | tgtcacctttcactccttAttctccttggagtcctttatc |
| Q580X | gtttacctggcactcctggaTaggatggattgccagggc | gccctggcaatccatcctAtccaggagtgccaggtaaac |
| Q700X | acctggtagcaaaggaTaaccaggtatccctggaattg | caattccagggatacctggttAtcctttgctaccaggt |
| L755X | gaaccaggatttgcatGacctgggccacct | aggtggcccaggtCatgcaaatcctggttc |
| Q928X | cttaaaggtgatgatggcttgTagggtcagccaggacttc | gaagtcctggctgaccctAcaagccatcatcacctttaag |
| Q930X | gatgatggcttgcagggtTagccaggacttcctggcccta | tagggccaggaagtcctggctAaccctgcaagccatcatc |
| E989X | gacttcctggccctacaggaTaaaaaggtagtaaaggagagcc | ggctctcctttactacctttttAtcctgtagggccaggaagtc |
| Q1016X | gtaaccctggtctccctggaTagccaggtcttataggacct | aggtcctataagacctggctAtccagggagaccagggttac |
| Q1052X | gaccttctggagttcctggaTaacctggctccccaggattac | gtaatcctggggagccaggttAtccaggaactccagaaggtc |
| Q1061X | ctccccaggattacctggaTagaaaggcgacaaaggtgatc | gatcacctttgtcgcctttctAtccaggtaatcctggggag |
| S1071X | gtgatcctggtatttGaagcattggtcttccag | ctggaagaccaatgcttCaaataccaggatcac |
| K1097X | ccagggaaccctggtatcTaaggttctgtgggagatc | gatctcccacagaaccttAgataccagggttccctgg |
| Q1180X | gtattcctggaccagctggaTagaagggtgaaccaggtcaac | gttgacctggttcacccttctAtccagctggtccaggaatac |
| Q1234X | cacggtttccctggtgtgTagggtcccccaggcc | ggcctgggggaccctAcacaccagggaaaccgtg |
| K1320X | ggccgggtctcaatggaatgTaaggagatcctggtctcc | ggagaccaggatctccttAcattccattgagacccggcc |
| Q1383X | cctccaggaatccctggcTagcctgggctaaagggtc | gaccctttagcccaggctAgccagggattcctggagg |
| Q1499X | ggcttttctctcctgtatgtaTaaggaaataaaagagcccacgg | ccgtgggctcttttatttccttAtacatacaggagagaaaagcc |
| Q1501X | aataaaagagcccacggtTaagacttggggacggctg | cagccgtccccaagtcttAaccgtgggctcttttatt |
| C1521X | accatgcctttcatgttctgAaacatcaataatgtttgc | gcaaacattattgatgttTcagaacatgaaaggcatggt |
| Y1543X | caagaaatgactattcttaAtggctctctaccccagagccc | gggctctggggtagagagccaTtaagaatagtcatttcttg |
| W1538X | gaaatgactattcttactgActctctaccccagagc | gctctggggtagagagTcagtaagaatagtcatttc |
| R1563X | gcatccagccattcattagtTgatgtgcagtatgtgaagctcc | ggagcttcacatactgcacatcAactaatgaatggctggatgc |
| C1567X | cattagtcgatgtgcagtatgAgaagctccagctgtggtgatc | gatcaccacagctggagcttcTcatactgcacatcgactaatg |
| E1574X | agtcgatgtgcagtatgtTaagctccagctgtggtgatc | gatcaccacagctggagcttAacatactgcacatcgact |
| W1594X | ggatgggattctctgtgAattggttattccttca | tgaaggaataaccaatTcacagagaatcccatcc |
| Y1597X | ttctctgtggattggttaAtccttcatgatgcatacaag | cttgtatgcatcatgaaggaTtaaccaatccacagagaa |
| S1632X | cctgcttggaagagtttcgttGagctcccttcatcgaatg | cattcgatgaagggagctCaacgaaactcttccaagcagg |
| S1661X | gctggcaactgtagatgtgtAagacatgttcagtaaacctcag | ctgaggtttactgaacatgtctTacacatctacagttgccagc |
| R1674X | gcaggagacttgaggacaTgaattagccgatgtcaag | cttgacatcggctaattcAtgtcctcaagtctcctgc |
| R1683X | ggacacgaattagcTgatgtcaagtgtgcatg | catgcacacttgacatcAgctaattcgtgtcc |
| C1684X | gaggacacgaattagccgatgAcaagtgtgcatgaagagg | cctcttcatgcacacttgTcatcggctaattcgtgtcctc |
| Q1685X | ggacacgaattagccgatgtTaagtgtgcatgaagaggac | gtcctcttcatgcacacttAacatcggctaattcgtgtcc |
| K1689X | gccgatgtcaagtgtgcatgTagaggacattctcgagcggtg | caccgctcgagaatgtcctctAcatgcacacttgacatcggc |
| G869R | gtccagggatccccAgagcacctggtcctatagg | cctataggaccaggtgctcTggggatccctggac |

| | | |
|---|---|---|
| C29R | gcagaggctgcggctCgAtatgggtgttctccagg | cctggagaacacccataTcGagccgcagcctctgc |
| C29W | gcagaggctgcggcttgGtatgggtgttctccagg | cctggagaacacccataCcaagccgcagcctctgc |
| S36R | ctatgggtgttctccaggaCGaaagtgtgactgcagtggc | gccactgcagtcacactttCGtcctggagaacacccatag |
| S36C | ctatgggtgttctccaggatGCaagtgtgactgcagtggc | gccactgcagtcacacttGCatcctggagaacacccatag |
| S36W | ctatgggtgttctccaggatGGaagtgtgactgcagtggc | gccactgcagtcacacttCCatcctggagaacacccatag |
| E130Q | gatgcaatggaaccaagggaCaacgtggatttccaggcag | ctgcctggaaatccacgttGtcccttggttccattgcatc |
| C1521R | accatgcctttcatgttcCgAaacatcaataatgtttgc | gcaaacattattgatgttTcGgaacatgaaaggcatggt |
| C1521W | accatgcctttcatgttctgGaacatcaataatgtttgc | gcaaacattattgatgttCcagaacatgaaaggcatggt |
| R1563C | gcatccagccattcattagtGCatgtgcagtatgtgaagctcc | ggagcttcacatactgcacatGCactaatgaatggctggatgc |
| R1563W | gcatccagccattcattagtGgatgtgcagtatgtgaagctcc | ggagcttcacatactgcacatcCactaatgaatggctggatgc |
| C1567R | cattagtcgatgtgcagtaCgAgaagctccagctgtggtgatc | gatcaccacagctggagcttcTcGtactgcacatcgactaatg |
| C1567W | cattagtcgatgtgcagtatgGgaagctccagctgtggtgatc | gatcaccacagctggagcttcCcatactgcacatcgactaatg |
| W1594R | ggatgggattctctgCgAattggttattccttca | tgaaggaataaccaatTcGcagagaatcccatcc |
| W1594C | ggatgggattctctgtgCattggttattccttca | tgaaggaataaccaatGcacagagaatcccatcc |
| S1632R | cctgcttggaagagtttcgtCGagctcccttcatcgaatg | cattcgatgaagggagctCGacgaaactcttccaagcagg |
| S1632C | cctgcttggaagagtttcgttGCgctcccttcatcgaatg | cattcgatgaagggagcGCaacgaaactcttccaagcagg |
| S1632W | cctgcttggaagagtttcgttGGgctcccttcatcgaatg | cattcgatgaagggagcCCaacgaaactcttccaagcagg |
| R1683C | gaggacacgaattagcTgCtgtcaagtgtgcatgaag | cttcatgcacacttgacaGcAgctaattcgtgtcctc |
| R1683W | gaggacacgaattagcTgGtgtcaagtgtgcatgaag | cttcatgcacacttgacaCcAgctaattcgtgtcctc |
| C1684R | gaggacacgaattagccgaCgAcaagtgtgcatgaagagg | cctcttcatgcacacttgTcGtcggctaattcgtgtcctc |
| C1684W | gaggacacgaattagccgatgGcaagtgtgcatgaagagg | cctcttcatgcacacttgCcatcggctaattcgtgtcctc |
| K1689Y | gccgatgtcaagtgtgcatgTaCaggacattctcgagcggtg | caccgctcgagaatgtcctGtAcatgcacacttgacatcggc |
| K1689Q | gccgatgtcaagtgtgcatgCagaggacattctcgagcggtg | caccgctcgagaatgtcctctGcatgcacacttgacatcggc |
| C29R-2 (N term tag) | gtgcgatcgctgcggctCgAtatgggtgttctccaggatc | gtgcgatcgctgcggcTcGatatgggtgttctccaggatc |
| C29W-2 (N term tag) | gtgcgatcgctgcggcttGgtatgggtgttctccaggatc | gatcctggagaacacccatacCaagccgcagcgatcgcac |

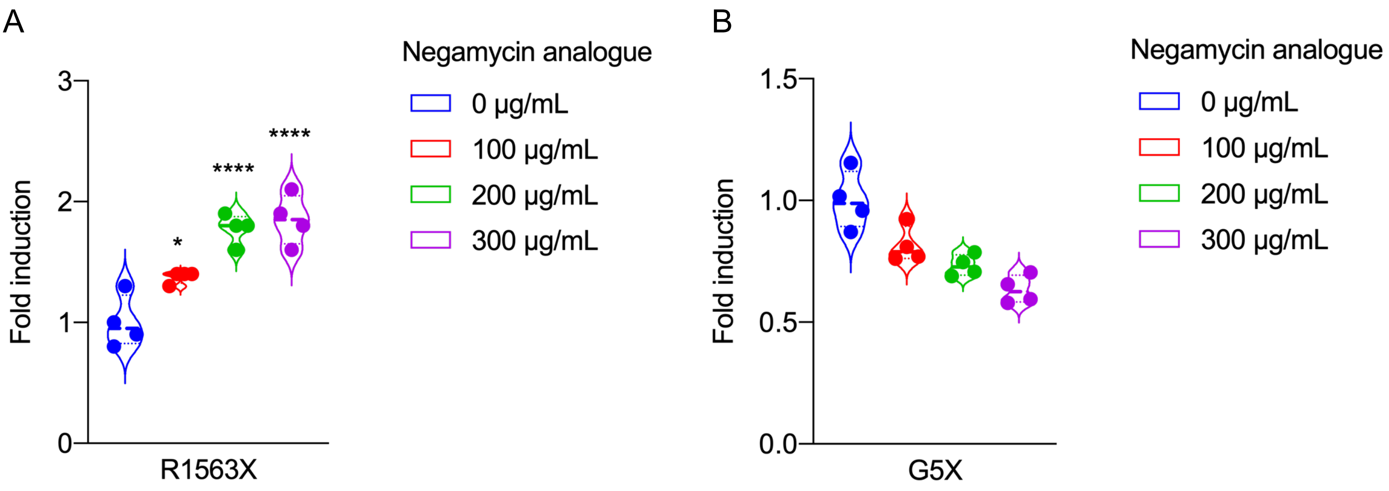


**Figure S1. Negamycin analogue CDX008 induces PTC-RT in the G418-susceptible COL4A5-R1563X mutant.** *A,B,* Luminescence was measured in cell lysates from HEK293 cells co-transfected with CMV-NanoLuc-fused COL4A5-R1563X plasmid and HSV-TK-Luc2 (firefly) for normalization. COL4A5-R1563X and -G5X-NanoLuc expressing cells were treated with serial dilutions of the indicated drugs. Negamycin analogue CDX008 significantly induced PTC-RT of COL4A5-R1563X.

Statistical analysis was performed using one-way ANOVA with Dunnett's multiple comparisons test (n=4). *, P <0.05; ****, P <0.001 vs. no treatment.

**G418 susceptible UGA**

| | -20 | -19 | -18 | -17 | -16 | -15 | -14 | -13 | -12 | -11 | -10 | -9 | -8 | -7 | -6 | -5 | -4 | -3 | -2 | -1 | PTC | 1 | 2 | 3 | 4 | 5 | 6 | 7 | 8 | 9 | 10 | 11 | 12 | 13 | 14 | 15 | 16 | 17 | 18 | 19 | 20 |
|---|---|---|---|---|---|---|---|---|---|---|---|---|---|---|---|---|---|---|---|---|---|---|---|---|---|---|---|---|---|---|---|---|---|---|---|---|---|---|---|---|---|
| C29X | a | g | c | c | t | g | c | a | g | a | g | g | c | t | g | c | g | g | c | t | t g A | t | a | t | g | g | g | t | g | t | t | c | t | c | c | a | g | g | a | t | c |
| S36X | g | c | t | a | t | g | g | g | t | g | t | t | c | t | c | c | a | g | g | a | t G a | a | a | g | t | g | t | g | a | c | t | g | c | a | g | t | g | g | c | a | t |
| C1521X | g | t | a | c | c | a | t | g | c | c | t | t | t | c | a | t | g | t | t | c | t g A | a | a | c | a | t | c | a | a | t | a | a | t | g | t | t | t | g | c | a | a |
| R1563X | g | c | a | t | c | c | a | g | c | c | a | t | t | c | a | t | t | a | g | t | T g a | t | g | t | g | c | a | g | t | a | t | g | t | g | a | a | g | c | t | c | c |
| C1567X | t | c | a | t | t | a | g | t | c | g | a | t | g | t | g | c | a | g | t | a | t g A | g | a | a | g | c | t | c | c | a | g | c | t | g | t | g | g | t | g | a | t |
| W1594X | c | t | c | a | g | g | g | a | t | g | g | g | a | t | t | c | t | c | t | g | t g A | a | t | t | g | g | t | t | a | t | t | c | c | t | t | c | a | t | g | a | t |
| S1632X | c | c | t | g | c | t | t | g | g | a | a | g | a | g | t | t | t | c | g | t | t G a | g | c | t | c | c | c | t | t | c | a | t | c | g | a | a | t | g | t | c | a |
| R1683X | a | c | t | t | g | a | g | g | a | c | a | c | g | a | a | t | t | a | g | c | T g a | t | g | t | c | a | a | g | t | g | t | g | c | a | t | g | a | a | g | a | g |
| C1684X | t | g | a | g | g | a | c | a | c | g | a | a | t | t | a | g | c | c | g | a | t g A | c | a | a | g | t | g | t | g | c | a | t | g | a | a | g | a | g | g | a | c |

**G418 non-susceptible UGA**

| | -20 | -19 | -18 | -17 | -16 | -15 | -14 | -13 | -12 | -11 | -10 | -9 | -8 | -7 | -6 | -5 | -4 | -3 | -2 | -1 | PTC | 1 | 2 | 3 | 4 | 5 | 6 | 7 | 8 | 9 | 10 | 11 | 12 | 13 | 14 | 15 | 16 | 17 | 18 | 19 | 20 |
|---|---|---|---|---|---|---|---|---|---|---|---|---|---|---|---|---|---|---|---|---|---|---|---|---|---|---|---|---|---|---|---|---|---|---|---|---|---|---|---|---|---|
| G5X | t | c | t | a | g | a | g | c | a | t | g | a | a | a | c | t | g | c | g | t | T g a | g | t | c | a | g | c | c | t | g | g | c | t | g | c | c | g | g | c | t | t |
| R266X | a | t | c | a | g | g | g | a | c | t | t | c | c | t | g | g | t | g | a | c | T g a | g | g | g | c | c | t | c | c | t | g | g | a | c | c | t | c | c | a | g | g |
| R373X | t | g | c | c | t | g | g | a | g | a | a | a | a | a | g | g | a | g | a | g | T g a | g | g | a | t | t | t | c | c | t | g | g | a | a | t | a | c | a | g | g | g |
| L755X | a | g | g | g | t | g | a | a | c | c | a | g | g | a | t | t | t | g | c | a | t G a | c | c | t | g | g | g | c | c | a | c | c | t | g | g | g | c | c | a | c | c |
| S1071X | a | c | a | a | a | g | g | t | g | a | t | c | c | t | g | g | t | a | t | t | t G a | a | g | c | a | t | t | g | g | t | c | t | t | c | c | a | g | g | t | c | t |
| W1538X | c | a | a | g | a | a | a | t | g | a | c | t | a | t | t | c | t | t | a | c | t g A | c | t | c | t | c | t | a | c | c | c | c | a | g | a | g | c | c | c | a | t |
| S1661X | g | g | c | t | g | g | c | a | a | c | t | g | t | a | g | a | t | g | t | g | t G a | g | a | c | a | t | g | t | t | c | a | g | t | a | a | a | c | c | t | c | a |
| R1674X | a | a | g | c | a | g | g | a | g | a | c | t | t | g | a | g | g | a | c | a | T g a | a | t | t | a | g | c | c | g | a | t | g | t | c | a | a | g | t | g | t | g |

**G418 susceptible UAG, UAA**

| | -20 | -19 | -18 | -17 | -16 | -15 | -14 | -13 | -12 | -11 | -10 | -9 | -8 | -7 | -6 | -5 | -4 | -3 | -2 | -1 | PTC | 1 | 2 | 3 | 4 | 5 | 6 | 7 | 8 | 9 | 10 | 11 | 12 | 13 | 14 | 15 | 16 | 17 | 18 | 19 | 20 |
|---|---|---|---|---|---|---|---|---|---|---|---|---|---|---|---|---|---|---|---|---|---|---|---|---|---|---|---|---|---|---|---|---|---|---|---|---|---|---|---|---|---|
| E130X | g | a | t | g | c | a | a | t | g | g | a | a | c | c | a | a | g | g | g | a | T a a | c | g | t | g | g | a | t | t | t | c | c | a | g | g | c | a | g | t | c | c |
| K1689X | g | c | c | g | a | t | g | t | c | a | a | g | t | g | t | g | c | a | t | g | T a g | a | g | g | a | c | a | t | t | c | t | c | g | a | g | c | g | g | t | g | t |

**Figure S2. cDNA sequence surrounding PTCs in G418-susceptible and non-susceptible mutants**
